# Enhanced intrathecal drug dispersion achieved by high volume injection and natural micromixing

**DOI:** 10.1101/2022.03.22.485353

**Authors:** Ayankola O. Ayansiji, Daniel S. Gehrke, Bastien Baralle, Ariel Nozain, Meenesh R. Singh, Andreas A. Linninger

**Affiliations:** Department of Bioengineering, University of Illinois Chicago, Chicago, Illinois, USA; UIC student intern from EPF, Ecole D’Ingénieur, Paris, France; Department of Chemical Engineering, University of Illinois Chicago, Chicago, Illinois, USA; Department of Neurosurgery, University of Illinois Chicago, Chicago, Illinois, USA

**Author notes:** **Corresponding author:** Andreas A. Linninger, Department of Biomedical Engineering and Neurosurgery, University of Illinois at Chicago, Chicago, IL, USA.

**Keywords:** Geometry Induced Mixing, Intrathecal Drug Delivery, Method of Moments, Oscillatory Flow, In vitro Spine Model, Drug Infusion Parameters

## Abstract

**Background:** Traditionally, there is a widely held belief that drug dispersion after intrathecal (IT) delivery is confined locally near the injection site. We posit that high volume infusions can overcome this perceived limitation of IT administration.

**Methods:** To test our hypothesis, subject-specific deformable phantom models of the human central nervous system were manufactured so that tracer infusion could be realistically replicated in vitro over the entire physiological range of pulsating cerebrospinal fluid (CSF) amplitudes and frequencies. Dispersion of IT injected tracers was studied systematically with high-speed optical methods to determine the relative impact of injection parameters including infusion volume, flow rate, catheter configurations and natural CSF oscillations.

**Results:** Optical imaging analysis of high-volume infusion experiments showed that tracer spreads quickly throughout the spinal subarachnoid space (SAS), reaching the cervical region in less than ten minutes. The experimentally observed biodispersion is much faster than suggested by prior theories (Taylor-Aris-Watson TAW dispersion). Our experiments indicate that micro-mixing patterns induced by oscillatory CSF flow around microanatomical features such as nerve roots significantly accelerate solute transport. Strong micro mixing effects due to anatomical features in the spinal subarachnoid space were found to be active in intrathecal drug administration but were not considered in prior dispersion theories. Their omission explains why prior models developed in the engineering community are poor predictors for IT delivery.

**Conclusion:** Our experiments support the feasibility of targeting large sections of the neuroaxis or brain utilizing high-volume IT injection protocols. The experimental tracer dispersion profiles acquired with an in vitro human CNS analog informed a new predictive model of tracer dispersion as a function of physiological CSF pulsations and adjustable infusion parameters. The ability to predict spatiotemporal dispersion patterns is an essential prerequisite for exploring new indications of IT drug delivery that targets specific regions in the central nervous system (CNS) or the brain.

## Background

Early studies on intrathecal (IT) administration in pigs using very slow infusion rates [1] contributed to the widely held belief that IT administration is confined to a small location near the injection site and thus is unsuitable for drug targeting of the brain. This notion enjoys theoretical support from the conjecture that IT drug delivery follows the phenomenon of solute dispersion in oscillatory pipe flow, well-known as Taylor dispersion in the engineering community [2], [3]. However, the applicability of Taylor dispersion on drug transport in oscillatory CSF flow has not been tested in vivo due to technical difficulties and risk to patients. It would further be tempting to test whether the theory of solute transport in oscillatory baffled reactors (OBR) [4], [5] also applies to IT delivery. Tracking tracer dispersion in vivo with multimodal PET/MRI [6] or computed tomography angiography suffers from limitations in spatial and temporal feature resolution [7]. Especially, observation of fast injection jets would require real-time acquisition rates unavailable in current non-invasive imaging technology [8]. Hence, technical limitations for tracking solutes suspended in complex CSF flow and patient safety make in vivo quantification problematic, if not impractical. In vitro experiments using an anatomically accurate model of the spinal subarachnoid space (SAS) are a compliment and a logical alternative to authentic, but often inaccurate in vivo infusion trials with limited temporal and spatial resolution. Several labs have also employed in vitro models for studying CNS dynamics [9]–[12]. IT *bench top testing* with optical image analysis offers the distinct benefit of achieving high temporal and spatial resolution, which is necessary for systematic parameter studies of the correlation between infusion and physiological parameters (=anatomy and CSF dynamics) and achievable drug distributions. A key requirement for realistic infusion bench tests is the availability of an anatomically accurate model of the spinal microanatomy with a deformable, fluid-filled spinal compartment with controllable pulsatile CSF flow.

In this paper, we will present parametric studies of IT infusion experiments in a subject-specific, anatomically accurate, 3D printed, transparent replica of the human spinal subarachnoid spaces (SAS) with natural CSF pulsations within the physiological range of spinal fluid amplitude and frequency [13]. High-speed video recording enabled accurate observation of spatiotemporal tracer distribution patterns following high-volume IT injection as a function of natural CSF oscillations. The results characterize speed of tracer front (thereafter referred to as *dispersion speed*) as a function of infusion settings (infusion volume, flow rate, position, duration, catheter diameters) and natural physiological properties (i.e. CSF stroke volume amplitude and frequency). We further compare the experimental data with prior theories (Taylor, OBR) of solute transport in oscillatory pipe flow.

## Materials and Methods

### In vitro Human Spine and Central Nervous System Model

We designed a deformable model of the human central nervous system (CNS) to reproduce functional biomechanical relations between dynamically interacting CSF compartments (Figs. 1a, b). Anatomically accurate analogue of the spinal subarachnoid spaces (SAS) with the transparent spinal cord including pairs of peripheral nerve roots and the translucent dural surfaces were manufactured in a multistep 3D printing and casting process with subject-specific imaging data of a 26-year old male volunteer [13]. Spinal CSF motion was generated by transmitting oscillatory expansion and contraction of an inflatable balloon located in the head section to the fluid. The balloon, mimicking the cerebrocranial vascular bed, in turn was driven by a piston pump capable of generating stroke volumes up to 1ml/beat in the frequency range of 0-127 beats per minute, see details in Appendix A. The mode of pulsatile flow conditions in the spinal CSF-filled spaces of the bench model reproduces pulsatile vascular bed dilation consistent with our understanding of periodic intracranial CSF displacement [14]. More manufacturing details of the subject-specific CNS replica can be found elsewhere [15]–[17].

**Fig. 1.**
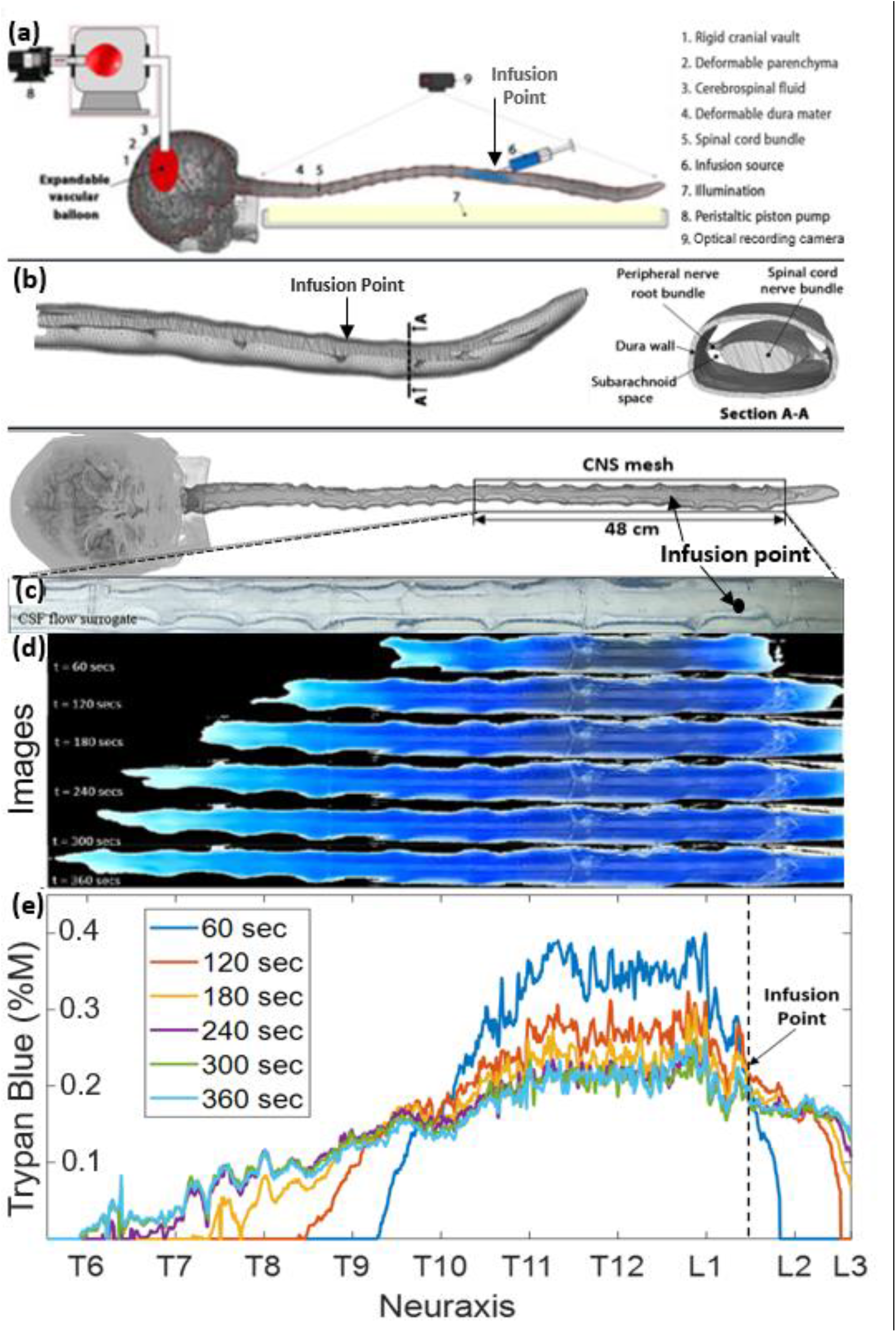
Phantom geometry, and videography of the dispersion of infused tracer in the central nervous system flow surrogate model. (a) A schematic diagram of the experimental setting. (b) the meshed model of a subject-specific central nervous system spine phantom models with peripheral nerve roots obtained from MR images. (c) Depiction of optically clear cerebrospinal flow surrogate model before the injection (no blue dye is visible yet). (d) Progressing tracer front spreading from the injection site preferably in cranial direction. CSF: cerebrospinal fluid. IVF: infusion volumetric flowrate. CNS: the central nervous system. (e) Overview of experiments and the time-lapsed images from a typical infusion experiment with IVF 2.0 ml/min, pulsation 1.0 ml/beat, and 0.4 %M of trypan blue concentration. It also shows a ruler with labels for the anatomical regions (i.e., cervical, thoracic, and lumbar) as well as the axial coordinate system, along the neuroaxis.

To reproduce conditions of in vivo IT procedures on the bench, infusion catheters with inner diameters of 0.2mm, 1.0mm, and 3.2mm were inserted into the lumbar and thoracic regions with inner diameters of 0.2 mm, 1.0 mm, and 3.2 mm. The elastic dura of the spine model has self-sealing property, thus enabling realistic catheter insertion and placement similar to clinical practice for human therapies. A wide range of infusion parameters settings (i.e. infusion volume, flow rate, position, duration) and systemic blood pump settings (i.e. stoke volume and frequency) enabled implementation of a comprehensive spectrum of infusion scenarios (bolus, chronic drug pump) occurring in the physiological range of CSF oscillations.. A total of 77 IT infusion experiments of trypan blue (Sigma, Aldrich) released with a programmable syringe pump (Harvard Instruments) were performed to precisely investigate the correlation between dye dispersion, CSF pulsations and infusion parameters. Trypan blue was chosen for its intense blue color needed in optical front tracking.

### Tracer dispersion tracking with videography

#### Automatic image processing

Snapshots were obtained from the experimental videos showing the dispersion of the tracer at different times. See Appendix B for more details. MATLAB 2019b was used for semi-automated image analysis and quantification of dye dispersion. A filing system stored key parameters for each experimental run: infusion volume, infusion molarity, infusion volumetric flow rate (IVF), experiment duration, subject orientation (supine), oscillation frequency, and oscillation amplitude. Each video frame captured red-green-blue (RGB) data in the range of 0 to 255 for each pixel at a location x and time. RGB values were converted to grayscale and concentration to track the expansion of the dispersion front of the trypan blue. See details in Appendix C. Analysis was divided into two phases: <u>Phase-1</u>, (acute infusion, t=0-1 min) covered the time the infusion pump actively discharged dye into the CNS replica. <u>Phase-2</u> (post infusion, t>1 min) further tracked dynamic tracer spread under the influence of natural CSF pulsations (without further infusion). The results display in this work are of those obtained using the grayscale method and not the binary method.

#### Optical analysis of tracer concentration

RGB triplet values for pixels along the neuraxis were recorded in each frame used for analysis (typically snapshots one minute apart). RGB color values were aggregated with a grayscale formula and intensities inferred from white-offset (white =no dye, blue = all dye). White-offset and area under the curve, AUC, scaling enabled quantitative dye concentration *C*(*x,t*), inference at a particular position, x, along the neuraxis, (see details in Appendix C).

#### Analysis of the dispersion front

A custom image analysis code was used to determine the speed and spatial position of the visible dye front. Figs. 1c, d depict a time-lapsed series of images showing the expansion of the blue tracer before and after the infusion at time points 0s, 60s, 120s 180s, 240s, 300s and 360s. Fig. 1e depicts tracer concentrations profiles inferred from extracted RGB pixel intensity as described in the Methods section.

#### Experimental determination of the speed of the advancement of the tracer front

The *method of moments* (MoM) was used to determine the apparent tracer dispersion velocity from video data, because it minimizes the sensitivity of optically acquired concentration profiles to uneven lighting conditions, scattering effects, and uncertainty in concentration inference from intensity data [18]. The MoM has been previously used to quantify dispersion in an annular tube [19]–[21]. We first determined the *speed of caudocranial motion* of the infusate by tracking shifts in the center of gravity of the tracer profile. The first moment *m*_1_(*t*) of the observed concentration profiles at time t is calculated as in Eq. (1), where *x* is the position along the neuroaxis, *x*_0_, marks the infusion point. The extreme limit of the ROI equal to full length from caudal to cranial aspects measured *x*_*m*_=48 cm. In each time frame, t, the first moment, *m*_1_(*t*), gives the location of the center of gravity, 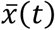 of the tracer concentration profile at a time point, *C*(*x,t*). The *speed of caudocranial motion* is equal to the change of the first moment with time.

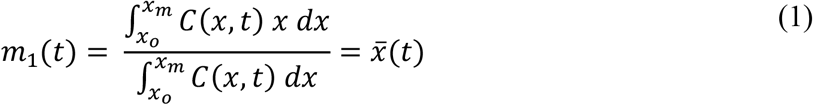

We further computed the second moment or *area moment of inertia* for each time point as in Eq. (2). The second moment can be interpreted as the mean spread of the visible concentration profile around its center, 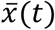. The increase in its variance with time, σ(*t*)^2^, is directly proportional to the apparent dispersion or diffusion rate *D*_*Ex*1_ of the tracer molecule as in Eq. (3). The coefficient of the apparent dispersion can then readily be determined as the rate of change in the second moment of the concentration curves by plotting their variance as a function of time. See Appendix D for details.

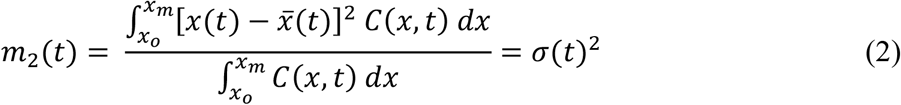

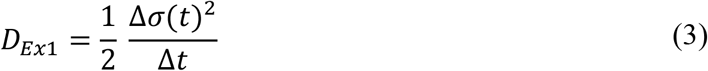

#### Stroke volumes and mean pulsatile flow velocities for formal analysis

In MR imaging, it is convenient to characterize natural CSF pulsations via the instantaneous total CSF volume, V(t), cervical stroke volume, *v*_c_, and CSF angular pulse frequency, *ω*. We estimated the average CSF flow velocity, U_rms_, with Eq. (4) to correlate clinical MR quantities to qu in formal flow analysis. Thus, two stroke volume settings of 0.5 to 1ml/beat in the frequency range of 0 to 120 beat/min enabled us to cover a wide range of CSF flow velocity averages covering the entire physiological range (0.09 cm/s – 0.93 cm/s).

The mean CSF flow velocity, *U*_*rms*_, in the cervical region was computed by integrating and averaging the squared ratio of pulsatile volumetric flow rate, 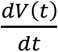, divided by the hydraulic cross-sectional area of the spinal CSF subarachnoid space, A, under cosine profile assumption as in Eq. (4), where V_0_ is the initial volume of CSF in cm^3^ in the system, N_bpm_ is the frequency in beat per minute, with period T in minutes.

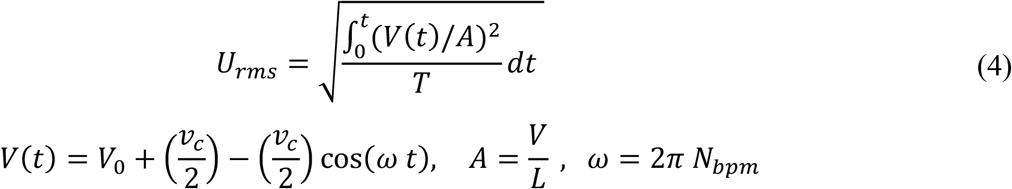

#### Statistical analysis

Initially, experiments were repeated three times with the same settings to ensure reproducibility and robustness of data acquisition. The results of the Shapiro-Wilk test indicated that all datasets had a normal distribution. ANOVA test was used in the regression analyses using IBM SPSS v26 to confirm that regression models described the dependent variables. Confidence intervals were calculated for all regression results. The Durbin-Watson statistic for all regression results was 0.716, therefore the datasets have autocorrelation. Furthermore, calculated tolerances indicate the multicollinearity effect in the datasets. A p-value of 0.05 was considered statistically significant.

## Results

### Dynamic tracer profile front tracking

Fig. 2a depicts tracer concentrations profiles inferred from calibrated RGB pixel intensity as described in the Methods section. Fig. 2b shows the first and the second moment for the experiment. The close agreement between three repetitions confirms reproducibility and acceptably low experimental variability of the experimental setting. The first moment gives information about the caudocranial velocity and the second moment gives information about the dispersion coefficient of the tracer as shown in Eq. (3), The binary mask was used to crop images to the width of the spinal section covered by detectable tracer intensity. The size of this mask along the neuraxis (x-direction) is referred to as the *dispersion width* of the tracer at time t, DD(t). The next section shows systematic analysis of infusion and physiological parameters on the speed of tracer dispersion.

**Fig. 2.**
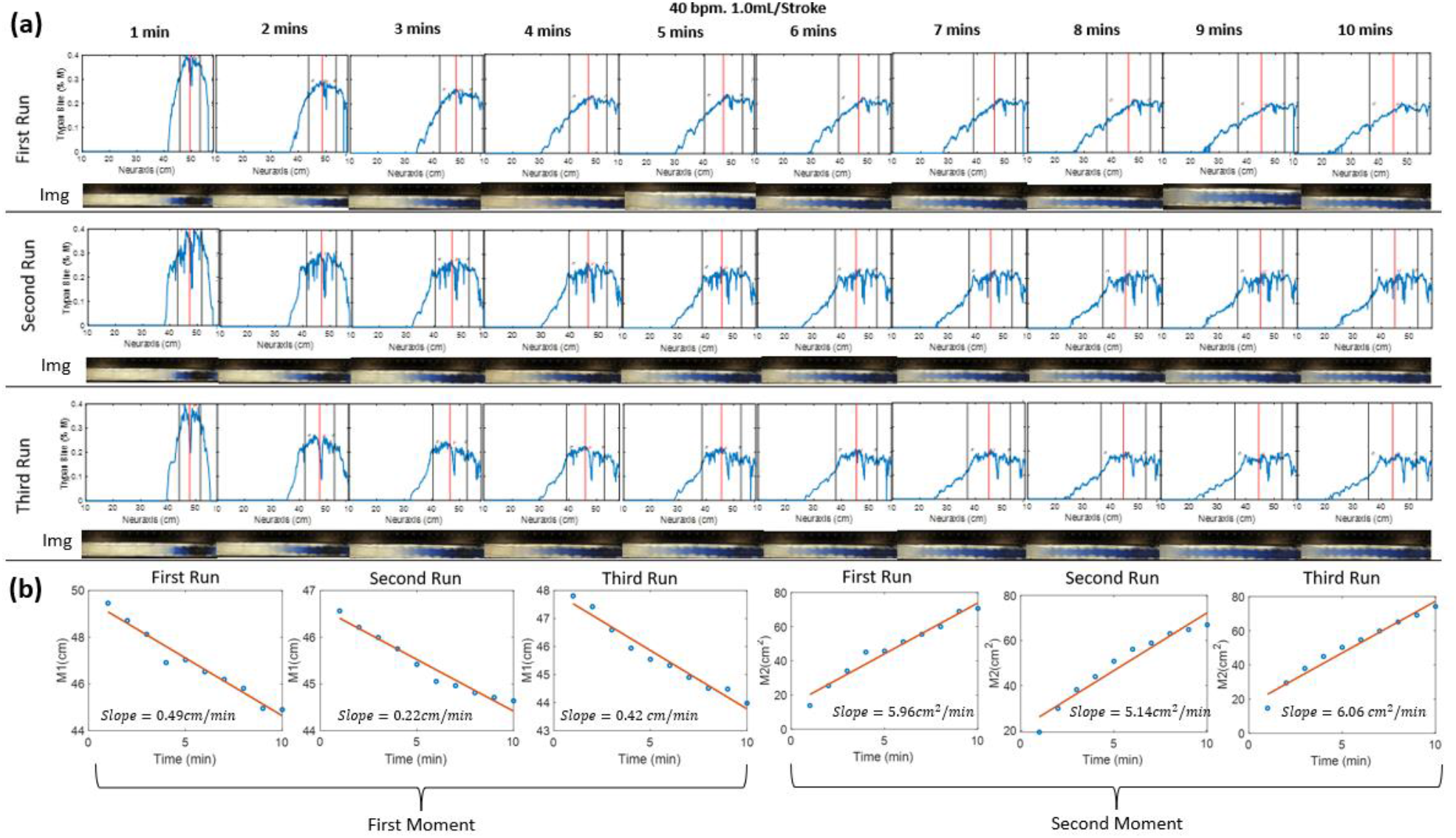
(a) The evolution of the tracer at different time of 40 bpm, 1.0mL/Stroke at different runs. Where the red and black vertical lines represent the mean and variance of the curve respectively. Img means image. (b) Shows the first and second moment for the three identical experiments. The small difference between repetitions suggest acceptable experimental variability. The first moment, M1, indicates that the tracer moves in cranial direction from the injection position (50 cm) towards the thoracic region position (44 cm). The second moments characterized the apparent speed of tracer dispersion. One half of the slope of second moment plot gives the dispersion coefficient

### Effect of catheter diameter on initial tracer spread (phase-1)

We studied the effect of catheter diameter on the speed and size of the dispersion front. Infusion experiments are performed using three different catheters with inner diameters of 0.2 mm (needle N1), 1.0 mm (N2), and 3.2 mm for widest needle (N3). Infusion lasted one minute over the course of 54 experiments. Dispersion width after 10 min, DD_10_ was measured from the point of the needle tip to the tip of the dye front in the caudal and cranial direction. The duration of 10 min was chosen, because this initial time window is critically important for assessing acute risks associated with high volume IT injection. High local toxicity has been implicated with granuloma formation [22], [23]. We also varied infusion volumetric flow rates (IVF= 0.5, 1.0, and 2.0 mL/min).

Fig. 3a shows that high caliber catheters promote slower tracer spread with shorter dispersion width in 1 min, DD_1_. For identical infusion flow rate (IVFs from 0.5, 1.0, and 2.0mL/min), the large caliber needle (N3) generated the shortest dispersion width. Fig. 3b depicts the effect of inner catheter diameter on the extent of the tracer spread observed 10 min after the infusion as a function of infusion flow. All experimental results (N=54) were also fitted into a linear regression model that can be used to estimate initial neuraxial coverage (=extent of the dispersion) as a function of catheter lumen and infusion volumes (IVF). The regression model of Eq. <u>(5)</u> can serve as a guideline in the clinical practice to estimate the length of initial dispersion as a function of catheter lumen and IVF (Fig. 3b).

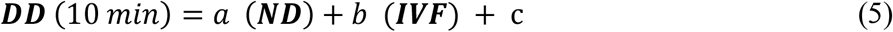

where *ND* is the inner needle diameter, and the *a, b, and c* are -1.4150, 2.8786, and 19.8668, respectively with R^2^ value of 0.7916.

**Fig. 3.**
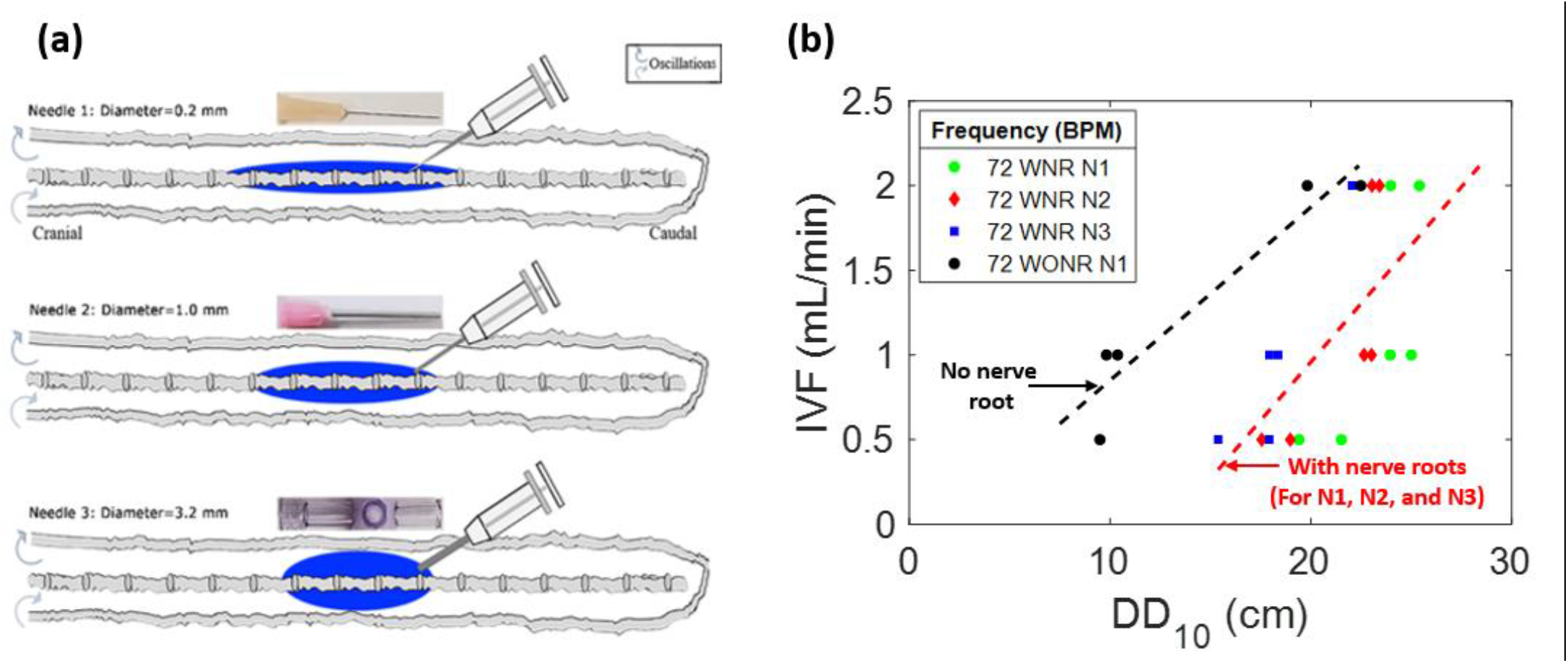
Effects of needle diameter, IVF, and DD on tracer dispersion. (a) The central nervous system (CNS) spine model is depicted as a diagram with varied diameter infusion catheters and corresponding idealized dispersion patterns (blue). (b) shows the relation of the infusion type to DD_10_ and needle diameter in a CNS spine model with peripheral nerve roots (colorful) and without peripheral nerve roots (gray). DD_10_ represents the linear dispersion distance the tracer front after 10 minutes of infusion. N1, N2, and N3 represent needles 1, 2, and 3, respectively. IVF: infusion volumetric flow rate. WNR: with nerve root. WONR: without nerve root.

The realization of a fixed infusion rate with thinner catheters requires higher infusion pressure resulting in higher exit velocities (*V*_*Ext*_) at their tips. Thus, catheter N1 at IVF=2.0 ml/min has the largest *V*_*Ext*_ and kinetic energy (*KE*) (As shown in Appendix A). Also, the exit velocity and kinetic energy of catheter N1 are higher than that of needles 2 and 3 with the same IVF. In thin catheters, a higher infusion impulse is delivered during the injection phase which translates into a wider dispersion length in the observed tracer front for the same injection flow rate.

### Effect of injection parameters on initial caudocranial dispersion (phase-1, acute infusion)

Experimental settings of injection flow rate, injection volume, and catheter specifications were varied to explore optimal conditions for targeting the cervical section or brain area. Fifty-four experiments (N=54) were conducted to systematically characterize tracer targeting towards the cranial compartment as a function of infusion parameters (phase-1, depicted in Fig. 4A). The results in Fig. 4b show that higher infusion flow rates IVF accelerate the speed of the tracer front advancing in the cranial direction from the infusion catheter tip. The distance the tracer front moves in the cranial direction from the infusion catheter tip for the CNS model in the supine position as observed 10 mins after the infusion is represented by S_DD._ This effect at high volume injections is due to the increased insertion kinetic energy (see Appendix A).

**Fig. 4.**
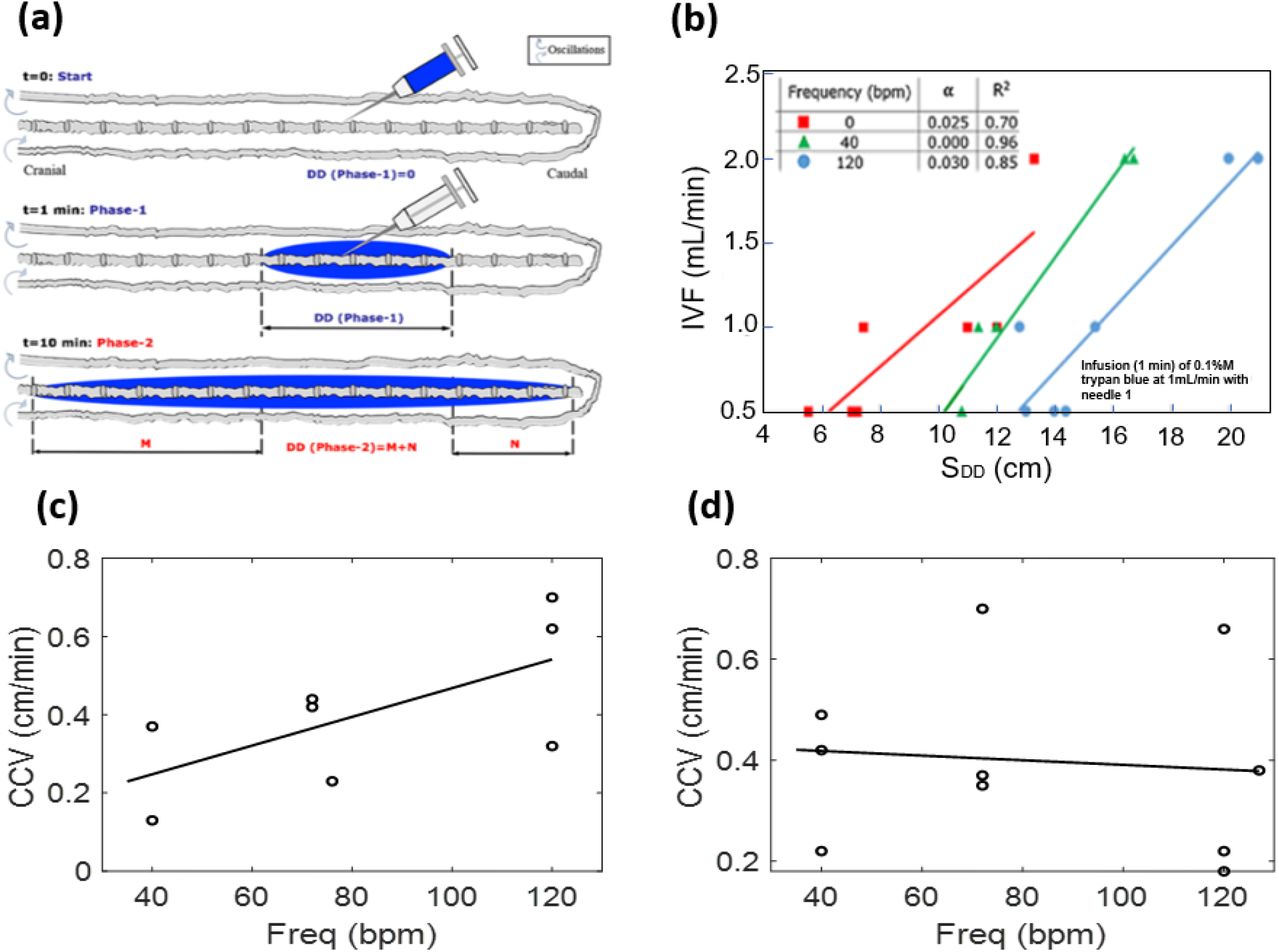
(a) Schematic depicting experimental setting in phase-1 tracer dispersion experiments. All results pertain to injection phase (phase-1) (b) Effect of CSF oscillation frequency on dispersion length S_DD_ (with nerves) using IVF of 0.5, 1.0, and 2.0 ml/min under oscillation frequencies 0, 40, and 140 bpm. (c, d) The caudocranial velocity (CCV) changes of the tracer relative to frequency for pulsation for 0.5 mL/beat (c) and 1.0 mL/beat (d) respectively. During the infusion phase, CCV is not frequency dependent, especially at higher CSF amplitudes.

The *speed of caudocranial motion*, CCV, is calculated as the change of the first moment with respect to time as shown in Fig. 2b. All experiments showed a shift of the first moment towards the cranium. We determined the velocity of apparent caudocranial advancement of the tracer front, ***CCV***, by recoding its positions for different time points. The results of Fig. 4C show that the caudocranial advancement of the tracer front from the injection site towards the cranium slightly increased with oscillation frequency, ***f***, as indicated by the regression of Eq. (6a) and *U*_***rms***_, as root-mean-square velocity, as indicated by the regression of Eq. (6b) with constant terms *a* and *b* as 0.3014 and 0.0012 for Eq. (6a), and 0.3752 and 0.0012 for Eq. (6b) respectively. At 1/ml stroke volume, this trend was similar but with a wider variability between runs (Fig. 4D)

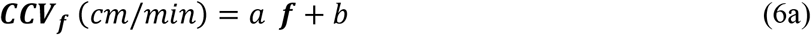

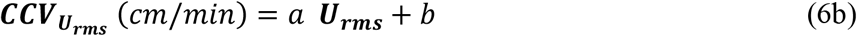

The rapid expansion of the drug front follows the amount of infused tracer qualitatively as expected, because the main driver of initial tracer spread is the injection impulse of fresh infusate concentrated in the relative narrow spatial confinement in the lumbar injection zone.

Eq (7) obtained by regression analysis describes the dependence of the effective diffusion coefficient on CSF pulse frequency for the phase 1. The terms *a* and *b* are constants that are obtained as 1.12 and 232.87 for stroke volume of 0.5mL/stroke and 3.62 and 232.63 for stroke volume of 1.0mL/stroke.

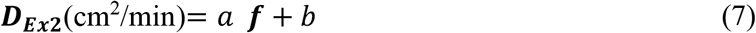

### Dispersion by natural CSF pulsation (phase-2, post injection)

Once the infusion stops (t >1min), further tracer spread is no longer propelled by injection impulse. It has long been known in the engineering community that oscillatory flow enhances solute dispersion, a phenomenon often referred to as Taylor-Aris dispersion (TAD). CSF flow spinal subarachnoid space is also oscillatory with zero net flux in our model. This is also approximately valid in vivo since bulk CSF production rates are much smaller than oscillatory fluxes.

The apparent dispersion coefficient of tracer spread in oscillatory CSF flow was determined experimentally (N=24) as a function of amplitudes (cervical CSF stroke volume, mean CSF flow velocities) and frequencies in a series of dynamic tracer infusion experiments. Two stroke volume settings of 0.5 to 1ml/beat in the frequency range of 0 to 120 beat/min enabled us to induce a wide range of CSF flow velocity averages covering the entire physiological range (0.09–0.93 cm/s). Tracer dispersion was rapidly reaching the cervical region in less than ten minutes, and it spread quickly throughout the spinal subarachnoid space (SAS). Experimentally obtained (=method of moments, MoM) dispersion coefficients, D_ex1_, summarized in Fig. 5 confirm our earlier finding [24], [25] that CSF amplitude and frequency are the critical factors of IT drug dispersion with the range of D_Ex1_=1.53 -2.94 cm^2^/min for a 0.5 mL/beat stroke volume, and D_Ex1_=2.57 – 4.70 cm^2^/min for the 1 mL/beat stroke volume as shown in Table E2 and E3, respectively, of Appendix E.

**Fig. 5.**
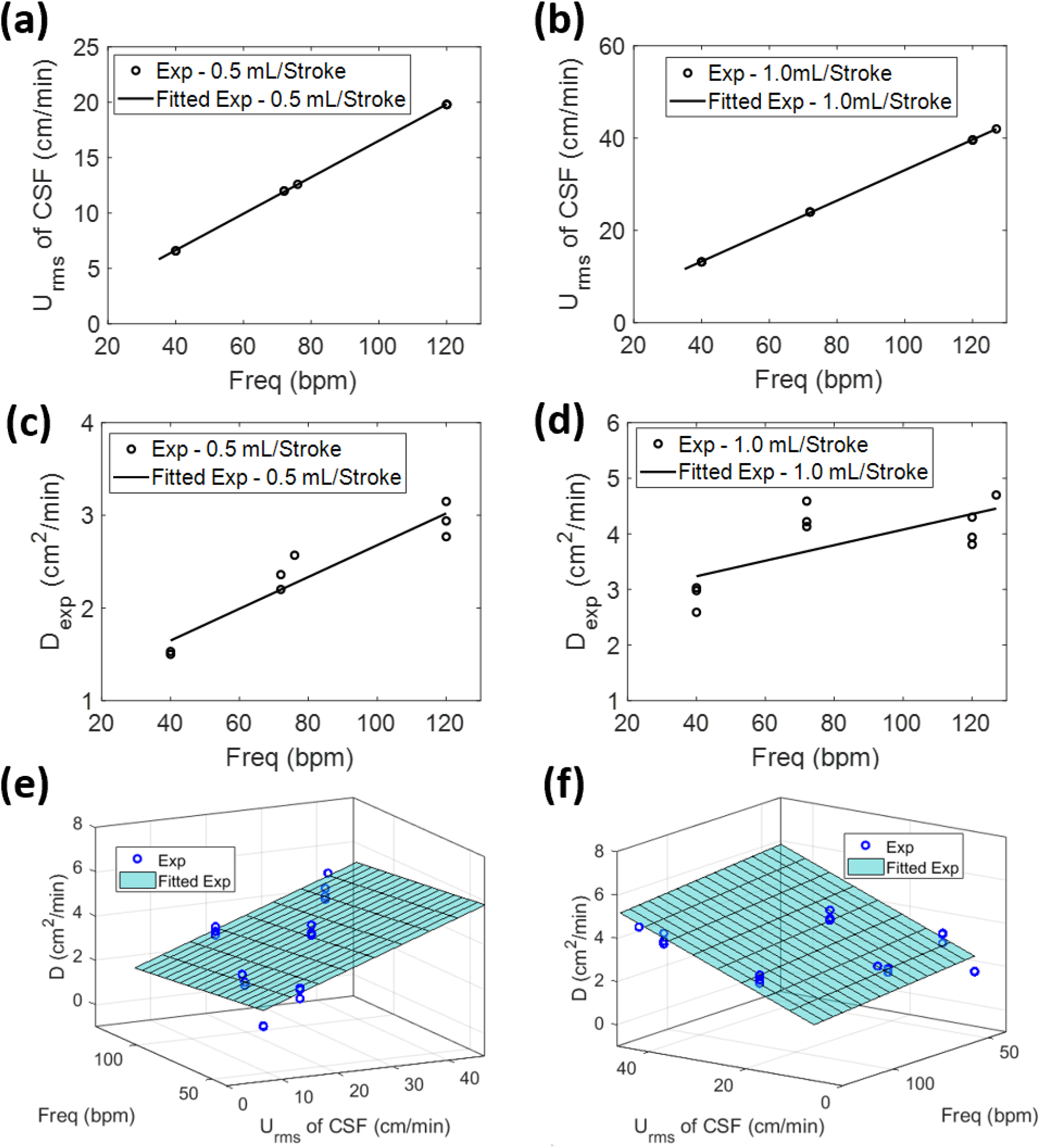
The root-mean-square velocity of the CSF at different frequencies for 0.5 mL/beat (a) and 1.0 mL/beat (b) respectively. Experimental diffusivity of infusion at different frequencies for 0.5 mL/beat (c) and 1.0 mL/beat (d). Relation of root-mean-square velocity of the CSF, frequency, and experimental diffusivity (e). (f) The same figure as (e) but shown from different perspective. In these results, 2 mL of 0.4 %M trypan blue was infused for 1.0 minutes as the infusion parameter. *D* is the calculated dispersion coefficient.

The correlation between apparent diffusion coefficient and CSF pulsations in Eq. (8), a function of CSF stroke volume was established using the Table E2 and Table E3 in Appendix E. It allows clinicians to estimate the effective dispersion in an intrathecal experiment as a function of the stroke volume in the cervical region, as well as the frequency of the CSF pulsations.

We also created a correlation between the apparent diffusion coefficient and CSF pulsations in Eq. (9a), a function of root-mean-square velocity of Eq. (9b). The mean CSF flow velocity, *U*_*rms*_, in the cervical region was computed as described in the methods section.

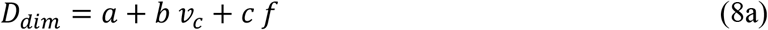

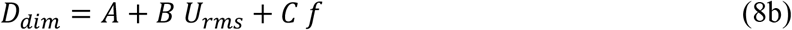

where a, b and c are constant with values -0.3390, 2.9051, and 0.0153 respectively. Also, where A, B and C are constants given as 1.9638, 0.0898, and -0.0083 respectively. *v*_*c*_ is the stroke volume, *U*_*rms*_ is the root-mean-square velocity of the CSF, *f* is the frequency and *D*_*dim*_ is the experimental dispersion coefficient for the dimensional model. This dimensional formula and the relationship supported by the data is given in Fig. 6A. The R^2^ value for Eq. (8b) is 0.74 and the variance is 0.93. Using this model (8b), the estimated dispersion coefficients are displayed in table E4 of Appendix E. The dimensionless form of Eq. (8b) was also developed where the dispersion coefficient is a function of Pe and Ω. As shown in (E7) of Appendix E

**Fig. 6.**
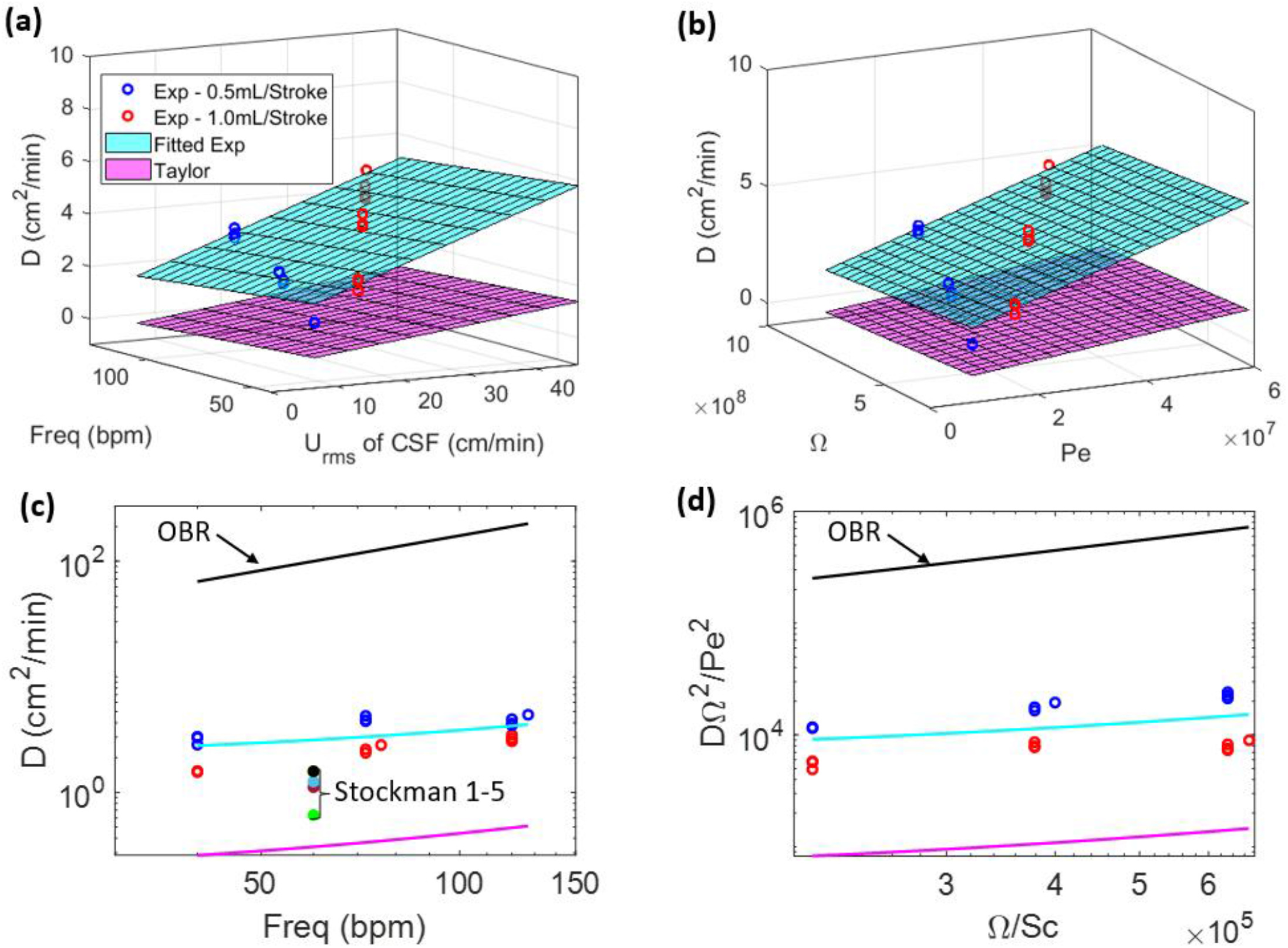
(a) The dispersion coefficients versus frequency at different root-mean-square velocity. The red and blue circles are experimental obtained dispersion coefficients at different frequencies and stroke volumes. (b) The dispersion coefficient as a function of the dimensionless numbers, Peclet number and dimensionless frequency (c) The comparison of the dispersion coefficient obtained experimentally, Taylor analysis, and the use of Oscillatory Baffle Reactor (OBR) for all stroke volumes. (i.e., 0.5 and 1.0 mL/Stroke) and different frequencies. The Stockman models were also used for frequency of 60bpm and stroke volume of 3.927e-6 mL/sec. (d)The dimensionless pairing of the dispersion coefficient with Peclet number and dimensionless frequency. Experimental values are from infusion experiment parameters of the following: 0.4% trypan blue infusion, IVF 2.0 mL/min, infusion volume 2.0 mL, supine orientation, with nerve roots, oscillation frequency of 34, 40, 72, 76, 118, and 120 bpm for 0.5 mL/beat and 40, 72, 86, 120, 127, and 169 bpm for 1.0 mL/beat. Where Stockman 1 – Stockman Spinal Model D, Stockman 2 – Stockman Model B-Normal, Stockman 3 – Stockman Model B-Dense, Stockman 4 – Stockman Model D-Normal, Stockman 5 – Stockman Model D-Dense.

### Effect of nerve roots on dispersion distance in phase-2

We observed previously that annular phantoms without microanatomical features underestimate actual dispersion after IT [26], [27]. To test the significance of microanatomical features on tracer dispersion, we also fabricated a spinal model without nerve roots and compared tracer propagation during infusion to the more anatomically realistic model with nerve roots. Tracer dispersion in the model with nerve roots was found to be always much more rapid than in a system lacking nerve roots under the same condition of oscillatory flow (Fig. 3b). Complex flow and mixing patterns observable are absent in idealized annular models lacking spinal microanatomy [3], [28], [29], thus failing to boost the effective dispersion of IT injected tracers as this is the case when microanatomical features are present. The results of this study provide further evidence for the significant impact of microanatomical features on the spatial and temporal dispersion patterns of IT administered solutes shown previously [13], [30].

### Is IT drug biodistribution predicted by Taylor-Aris Dispersion (TAD) theory?

Several authors [15], [29], [31] tried Taylor-Aris dispersion theory and its extension by Watson [32] to predict the intrathecal drug dispersion. Here we tested whether the Taylor-Aris-Watson (TAW) approach matches tracer dispersion in the bench infusion tests. Watson*’*s studies on solute dispersion in oscillatory pipe flow [32] based on prior work by Taylor [33] lead to analytical series solutions for the dispersion coefficient, D_3*D*_, in rectangular channels as a function of average velocity and frequency of the oscillatory flow field. Based on this work, Lee [6] reports the asymptotic limit of the dispersion coefficient D_3*D*_ of the solute in terms of group Schmidt, *Sc*, Peclet number, *Pe*, and dimensionless frequency, Ω, as in Eq. (9), where *U*_*rms*_ are the mean velocity, *ω* its oscillation frequency, *ν* the kinematic viscosity of the bulk fluid. The channel cross sectional dimensions are height and width *h* and*w*.

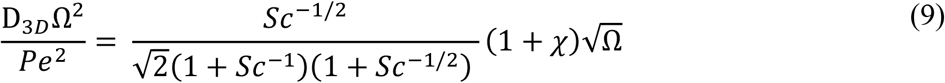

with

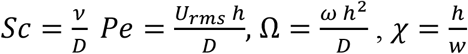

Actual dispersion coefficients and those predicted by the TAW as a function of the Peclet number and dimensionless frequency are shown in Fig. 6b. The mean CSF flow velocity, *U*_*rms*_, in the cervical region was computed as described in Eq. (4). It should be noted that in the CNS, the CSF velocity is graded along the neuraxis as a result of the spinal compliance[16] with the largest value in the cervical region and almost as zero in the sacral. In our analysis, we used the cervical *U*_*rms*_ for computing the Peclet number, so that the dispersion predicted by the TAD would be overestimated.

The data suggest that dispersion in IT experiments is much faster than predicted by TAD theory over the entire physiological range. Fig. 6a shows the relationship between the experimental and theoretical dispersion coefficients for stroke volume of 0.5 and 1.0mL/ beat. Figs. 6c-d shows the comparison of the dispersion coefficient obtained experimentally and using Taylor analysis for all the studied stroke volumes and frequencies. Also, the pioneering Monte Carlo simulation work by Stockman [34] on dispersion analysis in spinal subarachnoid space was also plotted. While the trends are correctly given, Stockman*’*s simulations predict about two times slower dispersion speeds than observed in our in vitro model. Fig. 6 also shows that the average dispersion coefficient obtained by the experiment is about 11 times bigger than that predicted by TAW.

### Reduced-order pharmacokinetic model for IT drug biodistribution

Experimental data for tracer infusion experiments served as input for a reduced order pharmacokinetic model of IT administration. The full description of this mechanistic drug administration model is beyond the scope of this manuscript but can be found elsewhere [35]. In brief, tracer biodispersion after lumbar intrathecal injection was simulated via a convection-diffusion process distributed along the neuraxis in Eq. (10). The effect of geometry induced mixing due to natural CSF pulsations was captured via the effective diffusivity, *D*_*eff*_, determined in experiments described in methods and results sections. Sensitivity of predictions to infusion settings was incorporated via a single source term, *V*_inj_(x,t). The effect of injection flow rate, volume and duration on the forces between infusate, spinal CSF interacting with deformable subarachnoid spaces (=dura) was accounted for by biomechanical fluid structure interaction (FSI). Accordingly, high volume injections generate a non-zero caudocranial convection term, *U*_*fsi*_, during infusion (phase-1). After the infusion stops (phase-2), tracer continues to spread away from the lumbar injection site in accordance with effective dispersion as in Eq. (10).

Tracer concentration profiles along the neuraxis as a function of time were predicted with mechanistic pharmacokinetic simulations. Pharmacokinetic tracer profile simulations with CNS dimensions and infusion settings used in the experiments took less than one CPU minute to converge generating asymmetrical profiles with peak concentration decreases with time (Fig. 7a). The comparison with the experimental run shows a qualitatively match both the spatial and the temporal dimension as shown in Fig. 7a. Fig. 7b shows maximum concentration is attained in a short interval of mean path length. The preliminary results of the prosed order model shows that the use of experimentally obtained effective dispersion coefficients can effectively predict drug dispersion after IT administration using a reduced order pharmacokinetic simulation. It is worth noting that simulation of biodistribution of active drugs into the CNS and the systemic circulation required additional information on biochemical parameters (denoted by the sink term *R*(*c, x*)) in Eq. (10) that might include drug half-life, tissue uptake and clearance[16].

**Fig. 7.**
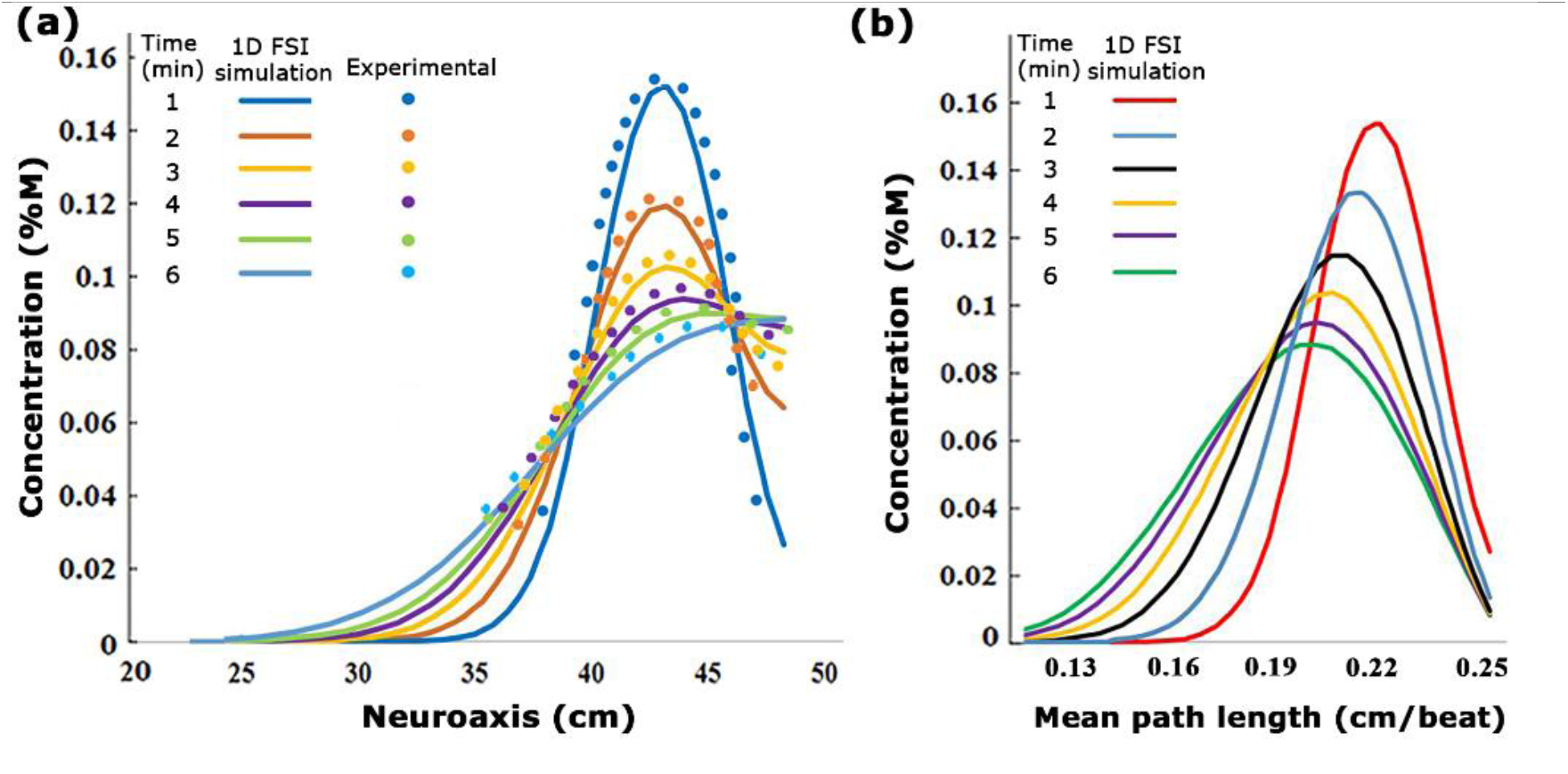
(a) A visualization of the experimentally derived concentration values compared to the 1D fluid-structure interaction (FSI) simulation concentration values at different times. (b) changes of 1D FSI simulation concentration diagrams in the different mean path lengths. The values are from infusion experiment parameters in 0.5 *mL*/*beat*.

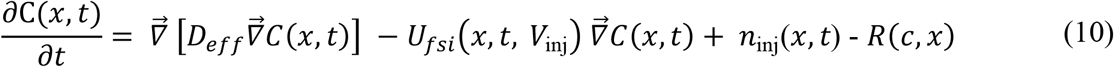

## Discussion

### Realistic replica of the human CNS

IT infusion studies under physiological conditions require deformability of the CSF spaces to accommodate realistic pulse pressure propagation and fluid motion inside a closed spinal SAS. Deformable fluid-filled spaces in the present bench test analog of the human CNS is fully enclosed between the soft dura/parenchyma surfaces and a distensible vascular interface (cranium vault with distensible vascular balloon) without leaving open system boundaries. The proposed configuration approximates the anatomy and fluid-structure interaction dynamics of the spinal SAS, so we have confidence that it reproduces the complex geometry induced CSF mixing patterns that were reported in previous work using direct numerical simulation of spinal CSF flow [13]. The CNS model also incorporates microanatomical features of a spinal cord, epidural space and peripheral nerve root bundles, especially, which are critical geometric aspects implicated with enhanced mixing [36], [37] of fluid layers in the spinal SAS leading to accelerated drug dispersion.

Moreover, transparent borders of the see-through human CNS replica enabled dynamic optical tracking of tracer concentration profiles during and after IT solute administration. Our experiments characterized in detail two stages occurring during high volume IT infusion: The initial phase during which the drug is injected lasts only a few minutes in clinical settings. The duration of 10 min was chosen, because it is suitable for assessing acute risks associated with high volume IT administration (local toxicity, granuloma formation).

### Injection phase

The axial dispersion during the injection phase (phase-1) correlated with infusion volume, IVF, and catheter diameters. Thinner catheters (inner diameter d=0.2 mm), generate wider and fast initial solute dispersion as is attainable with high caliber catheters (di=3.2 mm inner diameter). Based on our series of experiments, a simple formula in Eq (5) predicts initial dispersion width, DD_10_, the distribution length after 10 minutes of injection, as a function of injection volume/flow rate and catheter diameter. The formula in Eq. (5) may serve to estimate the initial volume of distribution, peak concentrations, and initial neuronal tissue exposure during the acute infusion phase of high-volume drug administration as a function of catheter lumen and infusion flow rate. Peak local toxicity risk may be elevated in infusion protocols generating narrower initial spread (i.e. high-volume infusion with large caliber catheters). It can also be used to estimate the expected local volume of drug action and local drug concentration to assess the risk of granuloma formations [23].

Under extremely low flow rate settings such as used in drug pumps, the effect of the injection impulse may be negligible. Accordingly, chronic administration with drug pumps may be governed by conditions of natural oscillation (phase-2) for its entire time course discussed below. For more recent developments for effective infusion protocols from drug pumps we refer to the work by Yaksh [38].

### Dispersion in naturally oscillating CSF (after the injection subsides)

After the injection ceases, the tracer spreads due to natural CSF pulsations. Oscillatory fluid flow around microanatomical features create geometry induced mixing pattern which breaks the laminar flow field by introducing eddies and vortices around nerve roots. Localized and interspersed eddies were observed in the flow directly upstream and downstream of cylindrical peripheral nerve root bundles suspended in the flow (see video in supplementary information). Moreover, trabeculae can substantially enhance this effect as reported in [13].

### Caudocranial motion

In all infusion experiments, slow but steady caudocranial advancement of tracer front from the lumbar injection site towards the cranium was observed. We offer two explanations for experimentally observed caudocranial transport.

First, lumbar injection divides the space available for the solute to disperse into a smaller distal volume containing the sacral compartment and a larger cranially facing domain stretching from the catheter tip to the thoracic, the cervical and the cranial SAS. We observed that tracers initially spread equally in both sections which is consistent with a diffusive process. Subsequently, the advancing dye fronts fill the closed sacral domain faster due to its smaller size. Once the closed sacral region is occupied, the tracer profile center of gravity begins shifting towards the head. Asymmetric caudocranial tracer profile develops due to a boundary effect which was confirmed with a mechanistic diffusive transport model. Systematic experiments with varying CSF conditions (amplitude and frequency) enabled the determination of a formula for the velocity of the caudocranial shift as a function of CSF pulsations and frequency as in Eq (6).

The graded biomechanical deformation profile of the spinal SAS responsible for CSF pulse attenuation is a second factor. CSF pulse amplitude was shown to diminish gradually from the cervical towards the sacral region [39]. The CSF pulsations and average oscillatory CSF velocities, U_rms_, become larger towards the cervical region compared to the lumbar region where the pulse amplitude is almost zero; thus, effective dispersion tends to become faster in the caudocranial direction.

The infusion experiments were conducted in a closed CNS model with no net CSF generation or removal. This supports the notion that bulk CSF flow or absorption is *not necessary* for caudocranial drug dispersion to occur. Rather, experiments suggest that caudocranial transport of infused solutes can simply result from the asymmetry of the CSF spaces and graded CSF pulse amplitudes.

Recently, an interesting analytical solution for drug dispersion in the SAS was developed by Lawrence et al. 2019 [15]. Predicted dispersion inside an idealized annular geometry without inclusion of microanatomical features (e.g., nerve roots) also found solute transport controlled by convection with negligible Taylor dispersion. Numerical simulation experiments within the cervical subarachnoid space [30] also showed that spinal cord nerve roots increases drug dispersion. These results confirmed independently our prior theoretical studies and are in agreement with new experimental results presented here.

### Derivation of clinical guidelines from in vitro experiments

Tracer dispersion after high volume injection was very rapid reaching the cervical region in less than ten minutes, and it spread quickly throughout the spinal subarachnoid space (SAS), much faster than predicted by the TAD model. The apparent dispersion coefficient was robustly determined experimentally as a function of CSF amplitudes and frequencies. An empirical correlation (Eq. 8) between apparent diffusion coefficient and CSF pulsations, a function of CSF amplitude and oscillation was established. This formula is of clinical interest to predict tracer dispersion for intrathecal drug administration.

We provide a simple guideline for estimating the volume of distribution of the drug during the injection phase as a function of catheter caliber and injection volume based on the experimental data and model in Eq. (5). Table 1 summarizes the expected size of the injection front (dispersion length after 10 min) from the lumbar injection site. It can also be used to get an idea about the advancement from the injection site towards the cranium, since the moving front of the tracer profile advances at least half of DD_10_.

**Table 1.** Guide for caudocranial motion, DD_10_ as a function of injection needle diameter (top row) and infusion volumetric flow rate (first column).

| caudocranial motion<br>after 10 min from<br>injection site (cm) | d=0.1 mm<sup>++</sup> | 0.2 mm | 0.5 mm | 1.0 mm | 1.5 mm | 3.2 mm |
| --- | --- | --- | --- | --- | --- | --- |
| 0.25 mL/min<sup>+</sup> | 20.49 | 20.34 | 19.92 | 19.21 | 18.51 | 16.10 |
| 0.50 mL/min | 21.20 | 20.06 | 20.63 | 19.92 | 19.22 | 16.82 |
| 1.00 mL/min | 22.62 | 22.48 | 22.05 | 21.35 | 20.64 | 18.24 |
| 2.00 mL/min | 25.46 | 25.32 | 24.90 | 24.19 | 23.48 | 21.08 |
| 2.50 mL/min | 26.88 | 26.74 | 26.32 | 25.61 | 24.90 | 22.50 |

<sup>+</sup>Infusion volumetric flowrate, <sup>++</sup>Inner diameter

For tracer dispersion in oscillatory CSF flow (phase-2), Eq (8a) quantifies effective dispersion as a function of stroke volume and pulse frequency. The Table 2 derived from these data, enables estimation of the effective dispersion coefficient for clinical applications. For drug molecules with different molecular diffusion coefficients (i.e. drugs with substantially different properties of our tracer), it may be used as a first approximation when no data are available or when its *Pe* number is in the same range as for trypan blue with a diffusion coefficient of D=1.938e-4 cm^2^/min.

**Table 2:**
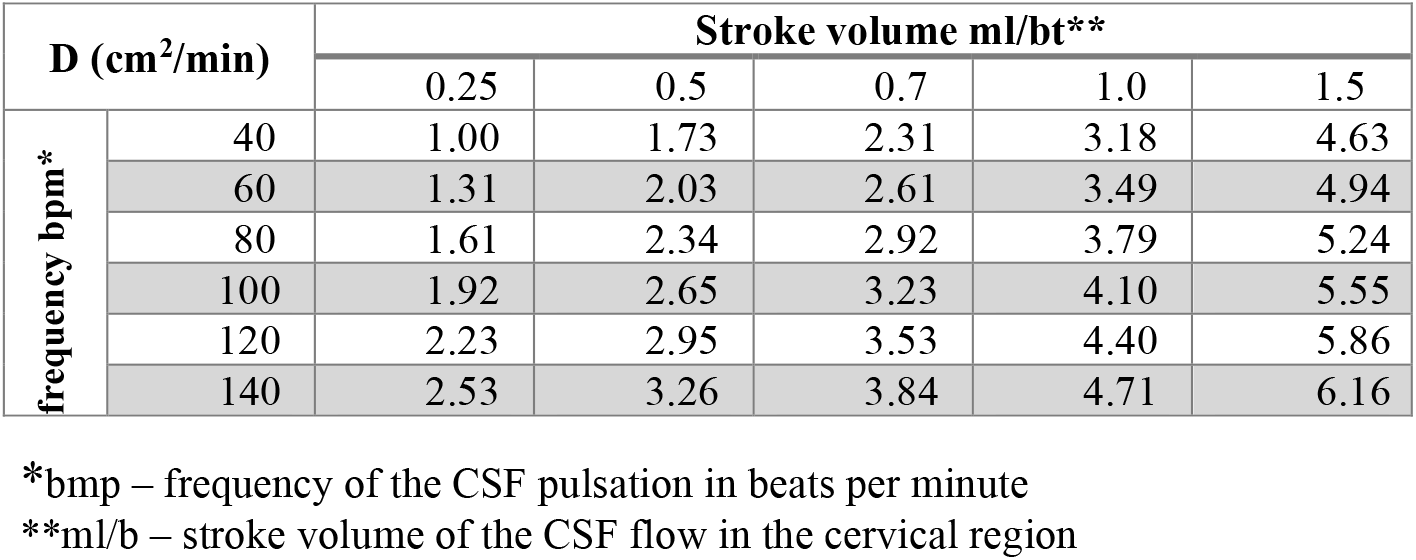
The table of clinical data, Dispersion Coefficient D

Note that the Schmidt number did not vary in our IT experiments, because CSF viscosity and molecular diffusivity of the tracer are constant. This is not a major restriction because drug dispersion typically operates typically in the high Schmidt number limit (in Eq. 9). where the authors [6] maintain that it is a very weak factor. Moreover, the viscosity of CSF is fixed in intrathecal drug delivery.

### Comparison to Taylor-Aris-Watson dispersion theory

The analytical Taylor-Aris-Watson approach was shown to underestimate the actual dispersion in vitro by about 12 times, despite our choice to use the maximum (=cervical) mean CSF velocity in the Peclet number. This is not entirely unexpected since several factors present in intrathecal drug dispersion do not meet the idealized assumptions in TAW approach. First, TAW does not account for geometry-induced mixing by micro-anatomical features. It also does not consider the effect of injection volume, catheter geometry and placement as discussed during phase-1 of the infusion experiments. Moreover, CSF flow with complex micro mixing occurs in a deformable spinal subarachnoid space, while TAW concerns rigid boundaries. Finally, CSF flow amplitudes are graded along the neuraxis, whereas TAW assumes constant flow velocity throughout the channel. We also quantified and compared prior work on baffled reactors (OBR) in the chemical engineering literature[4], [5] which provides further evidence in support of our main finding regarding the significance of geometry-induced mixing in solute dispersion in intrathecal drug administration. Appendix F shows that the Oscillatory Baffle Reactor theory over-estimated the dispersion coefficient by an average factor of 41.

### Eccentricity and stagnation zones

Several authors have implicated the eccentricity of idealized cross-sectional areas of CSF-filled spaces in the spinal subarachnoid space as a key factor for accelerated drug dispersion [31], [39], [40]. The insensitivity to centric or eccentric alignment in our experiments does not seem to support the notion of eccentricity as a significant factor for the speed of IT dispersion. Moreover, we could not observe stagnation or recirculation zones in any of the experiments.

### Limitations

There are safety limits to high volume injections adding the CSF amount during drug administration. We have experienced in rat that no more than 10% of the CSF volume can be safely injected over a period of a couple of minutes [41]. Higher injection impulse may also contribute to the possibility of high shear rates that nerve roots experience near the catheter tip, which may again pose an additional risk that requires clinical investigation.

The biomechanical stress-strain relation of the epidural spaces is a function of several poorly understood factors. These include viscous resistance exerted by venous blood volume and fatty tissues, possible elastic deformation resistance of nerve roots bulging into peripheral distal spaces, and the biomechanical properties of the dura membrane including the stiffening effect of by trabeculae and ligaments. The current model was able to induce cervical CSF displacements (stroke volume 0-1 ml/beat) within the physiological range but was not designed to faithfully reproduce the biomechanical compliance of the spinal compartment. Accordingly, we also did not attempt to measure absolute pressure changes that occur during infusion. We plan to study the rise in the line pressure or the increase in pressure in CSF of the spinal SAS as a function of injection parameters in future work.

Another related limitation pertains to the practice of fluid removal before high volume injection (initial CSF tapping). There was no attempt made to interrogate the CNS model regarding biomechanical response of the CSF spaces subject to tapping.

## Conclusions

We conducted an extensive parametric study of tracer dispersion in a subject-specific bench model of the human CNS which anatomical and functional reproduction of CSF dynamics from the spine to the cranium. A systematic variation of parameters in a large number of infusion experiments enabled in vitro quantification of the spatiotemporal tracer dispersion patterns and their dependence on significant infusion parameters and pulsatile CSF conditions. This study reports a unique set of experimental data on the combined effect of infusion and natural oscillations that have previously not been reported. Our bench experiments suggest the feasibility of targeting large sections of the neuroaxis to the brain with the help of high-volume injection protocols. Infusion studies using human CSN models may serve as an inexpensive surrogate for testing and optimizing infusion protocols for the safe distribution of IT administered solutes along the neuraxis to inform clinical trials in humans.

## Ethics approval and consent to participate

Not Applicable.

## Availability of data and material

The details of all data are available in the appendixes, tables, and figures.

## Author details

^1^Department of Biomedical Engineering, University of Illinois at Chicago, Chicago, Illinois, USA. ^2^Interning at UIC from EPF, Ecole D*’*Ingénieur, Paris, France. ^3^Department of Neurosurgery, University of Illinois at Chicago, Chicago, Illinois, USA.

## Acknowledgements

AL is grateful for funding from the National Science Foundation (https://www.nsf.gov), under grant number CBET-1301198. as well as the National Institute of Health, Neurological Disorders and Stroke (https://www.ninds.nih.gov) under Grant number NIH NINDS 1R21NS099896, and the National Institute of Aging (https://www.nia.nih.gov) under grant NIH NIA 1R56AG066634-01. We thank Dr. Gholampour at the University of Chicago for initial help with the statistical methods.

